# Taurine suppresses liquid-liquid phase separation of lysozyme protein

**DOI:** 10.1101/2021.01.26.428332

**Authors:** Kanae Tsubotani, Sayuri Maeyama, Shigeru Murakami, Stephen W Schaffer, Takashi Ito

## Abstract

Taurine is a compatible osmolyte that infers stability to proteins. Recent studies have revealed that liquid-liquid phase separation (LLPS) of proteins underlie the formation of membraneless organelles in cells. In the present study, we evaluated the role of taurine on LLPS of hen egg lysozyme. We demonstrated that taurine decreases the turbidity of the polyethylene glycol-induced crowding solution of lysozyme. We also demonstrated that taurine attenuates LLPS-dependent cloudiness of lysozyme solution with 0.5 or 1M NaCl at a critical temperature. Moreover, we observed that taurine inhibits LLPS formation of a heteroprotein mix solution of lysozyme and ovalbumin. These data indicate that taurine can modulate the formation of LLPS of proteins.

## Introduction

Recent studies have demonstrated that liquid-liquid phase separation (LLPS) of proteins and nucleic acids is associated with the assembly of membraneless organelle in cells. It plays a critical role in a variety of normal cellular activities; i.e. the formation of stress granules, nucleoli, and the condensation of proteins related to glucose metabolism and cell signaling (Banani et al. 2017; Kohnhorst et al. 2017; Zhang et al. 2020; Kamagata et al. 2020). Moreover, aberrant LLPS of many proteins, such as FUS, TDP-43, tau, and #alpha-synuclein, leads to fibril formation, which is related to neurodegenerative diseases (Zhang et al. 2017; Murray et al. 2017; Choi et al. 2018; Ray et al. 2020).

LLPS formation of proteins occurs through electrostatic interactions between amino acid residues, such as dipole-dipole, cation-π, #p-#p interactions (Nott et al. 2015; Vernon et al. 2018; Hofweber et al. 2018; Qamar et al. 2018). In many cases, LLPS formation requires a high protein concentration and changes in medium salt concentration, pH, temperature, pressure, etc (Zaslavsky et al. 2018; Cinar et al. 2019; Singh et al. 2020). In other cases, posttranslational modifications, such as phosphorylation and methylation, influence LLPS formation (Wippich et al. 2013; Wegmann et al. 2018; Ferreon et al. 2018; Milovanovic et al. 2018). Importantly, intrinsic small molecules also influence LLPS formation (Shiraki et al. 2020). For example, Patal et al. have demonstrated that ATP works as a hydrotrope, thereby inhibiting LLPS formation of proteins (Patel et al. 2017). This discovery raises the possibility of a novel role of cellular metabolites as regulators of LLPS formation.

Taurine is a beta-amino acid found in high concentration (typically 1∼40 mmol/kg tissue weight) in mammalian tissues (Chesney 1985; Ito et al. 2008; Jentsch 2016)v. Taurine functions as a compatible organic osmolyte, which regulates intracellular ionic balance, thereby maintaining cell volume (Schaffer et al. 2000, 2010). Compatible osmolytes also contribute to the thermodynamic stability of proteins by minimizing the water-protein interaction (Bruździak et al. 2018). The role of compatible osmolytes in LLPS formation of proteins has been evaluated by Cinar et al., who demonstrated that trimethylamine oxide (TMAO), which is an organic osmolyte of marine fish, enhances LLPS formation of gamma-crystallin(Cinar et al. 2019). Abe et al. demonstrated that taurine can weakly interact with hen egg lysozyme protein, including cation-#pai interaction, in crowding condition (Abe et al. 2015). Therefore, we hypothesized that taurine can regulate LLPS formation through its potential to interact with the protein surface, as well as directly with water.

Lysozyme is a frequently used model protein to study protein folding and aggregation. Lysozyme forms LLPS by adding a high concentration of salt or by mixing with ovalbumin (Taratuta et al. 1990; Muschol and Rosenberger 1997; Dumetz et al. 2008; Santos et al. 2018; Iwashita et al. 2018; Bye and Curtis 2019). Therefore, in the present study, we studied the effect of taurine on LLPS formation of lysozyme.

## Methods

### Materials

Hen egg lysozyme was kindly provided by QP Co. (Japan). Ovalbumin was obtained from Sigma-Aldrich (USA). Taurine and other chemicals were purchased from Nacalai Tesque (Japan).

### LLPS formation by high NaCl concentration

A standard Tris buffer [20 mM Tris-HCl (pH7.4)] was prepared. The pH of the Tris buffer was readjusted back to pH 7.4 after adding taurine. Lysozyme powder was dissolved in water and then mixed with standard Tris buffer and 5M NaCl solution. The lysozyme solutions were incubated at the indicated temperature (0∼20°C) by a Cool-thermo unit (Titec Co., Japan). After a 5 min incubation at each temperature, the cloudiness (whiteness) of the lysozyme solution was confirmed visually.

### LLPS formation by mixing lysozyme and ovalbumin

The formation of LLPS containing lysozyme and ovalbumin was performed according to previous reports (Santos et al. 2018; Iwashita et al. 2018), with a few modifications. In the first experiment, a lysozyme solution (#mg/mL in water) and an ovalbumin solution (#mg/mL in water) were mixed 1:1 ratio in water (final concentration is 2.5 mg/mL each). Taurine (0, 20, or 200 mM) was added before mixing the proteins. 20 #microL of the solution was applied to a disposal cell counter plate (Watson Co., Japan) and then LLPS formation was checked with optical microscopies (BZ-X800, Keyence, Japan). The turbidity of the solution was measured in 96-well microplates by using a UV-Vis microplate reader (Spectramax M2 microplate reader, Molecular Devices, USA).

In the latter experiments, lysozyme and ovalbumin were added to 5mM Tris-buffer (pH7.4). Then, LLPS formation and turbidity were analyzed. Additionally, FITC-labelled lysozyme (LS1-FC-1, Nanocs Inc., USA) instead of lysozyme was mixed with ovalbumin in 5mM Tris buffer (pH 7.4), and then LLPS formation was visualized by fluorescent microscopy (BZ-X800, Keyence).

## Results

### 1. Taurine inhibits the opalescence of lysozyme in supersaturated state

The crowding condition by adding polyethylene glycol (PEG) is a common method to observe LLPS of various proteins (Kaur et al. 2019). We tried to detect LLPS formation of lysozyme after crowding following addition of PEG-6000. Although the solution was cloudy, liquid droplet formation was not confirmed, as assessed by optical microscopy (Figure 1).

**Figure 1.**
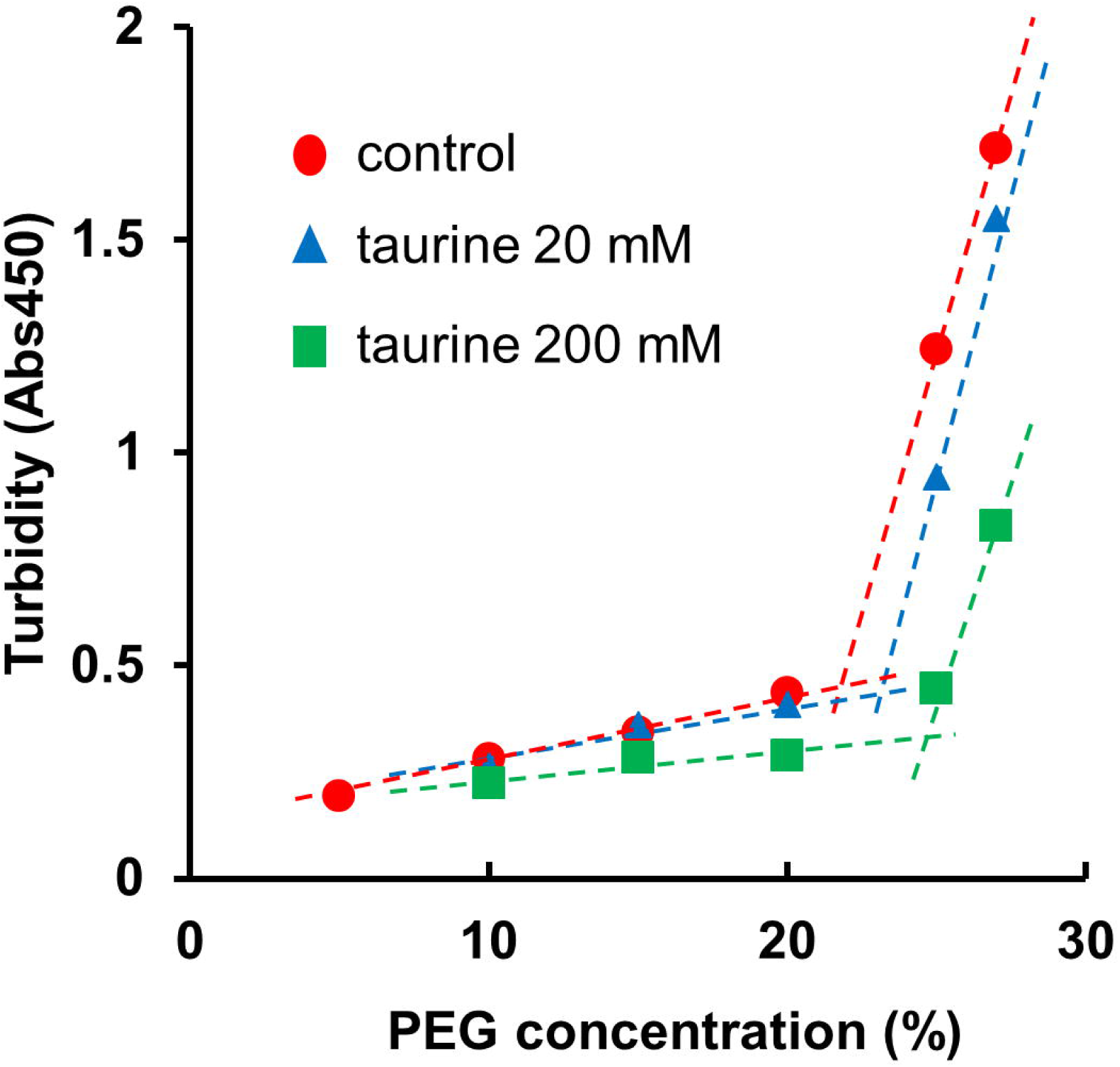
Absorbance (λ= 550nm) of PEG-containing lysozyme solution (50 mg). Effect of 20 and 100mM taurine on PEG-induced turbidity of lysozyme was measured.

We tested the effect of the cloudiness of the PEG-containing lysozyme solution. While increasing PEG concentration up to 20% increased the absorbance of the lysozyme solution (50 mg/mL) in Tris buffer (pH7.4) at room temperature, the absorbance was sharply increased at 25% PEG. The sharp increase in turbidity may be due to the precipitation of protein (Kaur et al. 2019). The addition of 100 mM taurine decreased the turbidity of the lysozyme solution (50 mg/mL). Figure 1B, C shows the turbidity data of PEG-containing lysozyme solution.

### 2. Taurine inhibits LLPS of lysozyme in the presence of a high NaCl concentration

Lysozyme forms LLPS in the presence of high NaCl and below a critical temperature (Muschol and Rosenberger 1997). To determine whether taurine can modulate LLPS formation, we tested the influence of taurine on the temperature at which LLPS started to form. Figure 2 shows the clouding point for lysozyme at two different NaCl concentrations, 0.5M, and 1 M, in the presence or absence of taurine (0, 20, 200 mM). The addition of taurine decreased the clouding point temperature for 100 mg/mL lysozyme in the presence of 0.5M NaCl; while the lysozyme solution without taurine started to become cloudy at 9 °C, the solution with 200mM taurine started to become cloudy at 6 °C while 20 and 50mM taurine shifted the clouding point to 8°C. In 1M NaCl solution, the addition of 200Mm taurine, but not 20 or 50mM, also decreased in the clouding point temperature; taurine shifted it from 26 °C to 24 °C in 100mg lysozyme solution.

**Figure 2.**
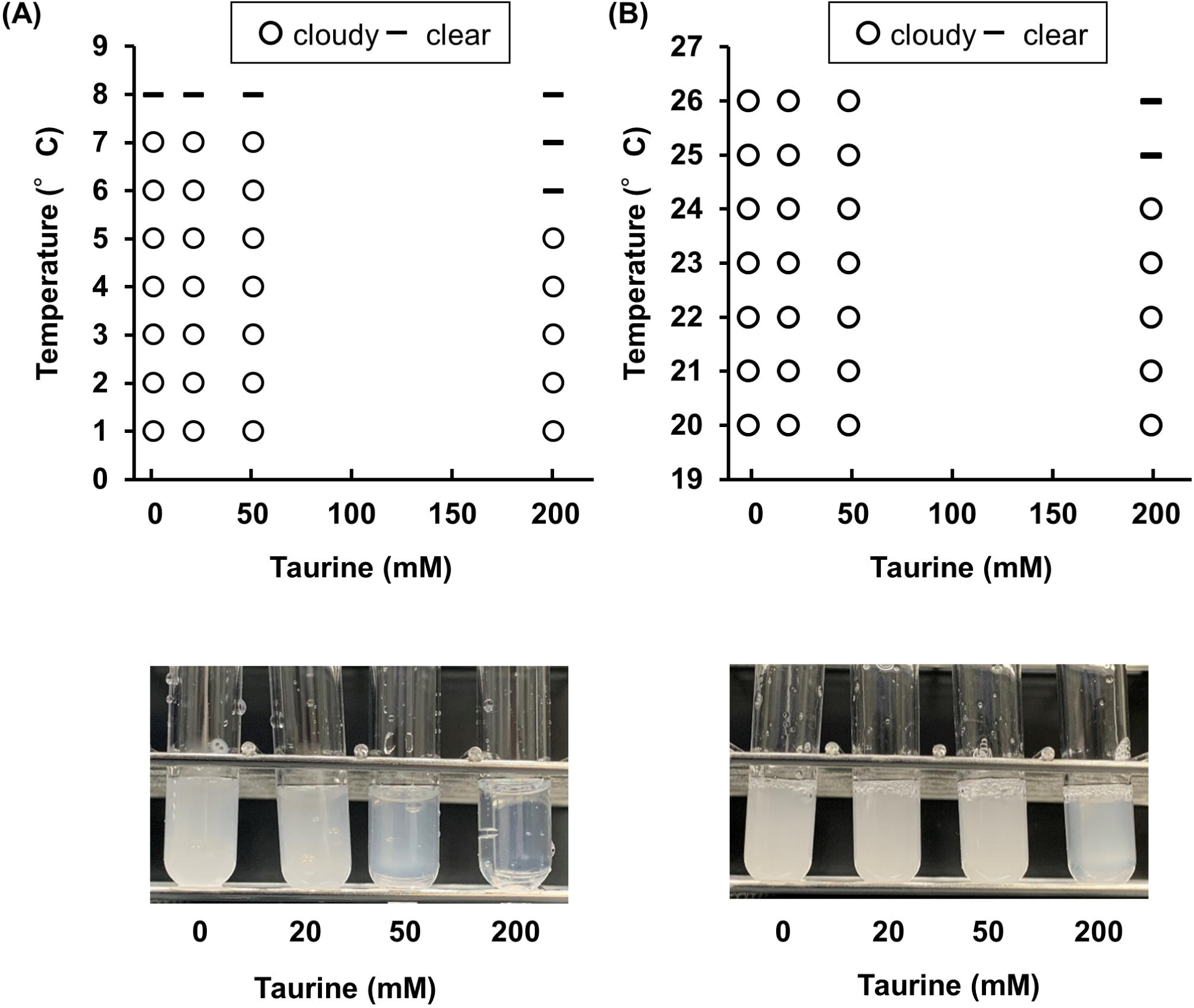
Effect of taurine on NaCl and temperature-dependent LLPS of lysozyme.(A,B) Cloud point data for lysozyme solution at 0.5M (A) or 1M (B) NaCl concentration with or without taurine. Lower picutres: epresentative clouded lysozyme solution in 0.5M NaCl at 9°C and in 1M at 26°C.

### 3. Taurine inhibits LLPS formation of heteroprotein mix, lysozyme and ovalbumin

Lysozyme forms LLPS by mixing with ovalbumin (Santos et al. 2018; Iwashita et al. 2018). Next, we tested the influence of taurine on heteroprotein LLPS formation of lysozyme and ovalbumin. LLPS formation was monitored by phase-contrast microscopy. Two proteins were mixed at a 1:1 ratio (1mg/mL each) in a Tris-buffered solution (5mM, pH7.4, Figure 3A). Mixing two proteins generates droplets. The addition of 20∼100 mM taurine decreased the size of droplets of the heteroprotein mix (Figure 3B). The addition of 200mM taurine prevented LLPS formation.

**Figure 3.**
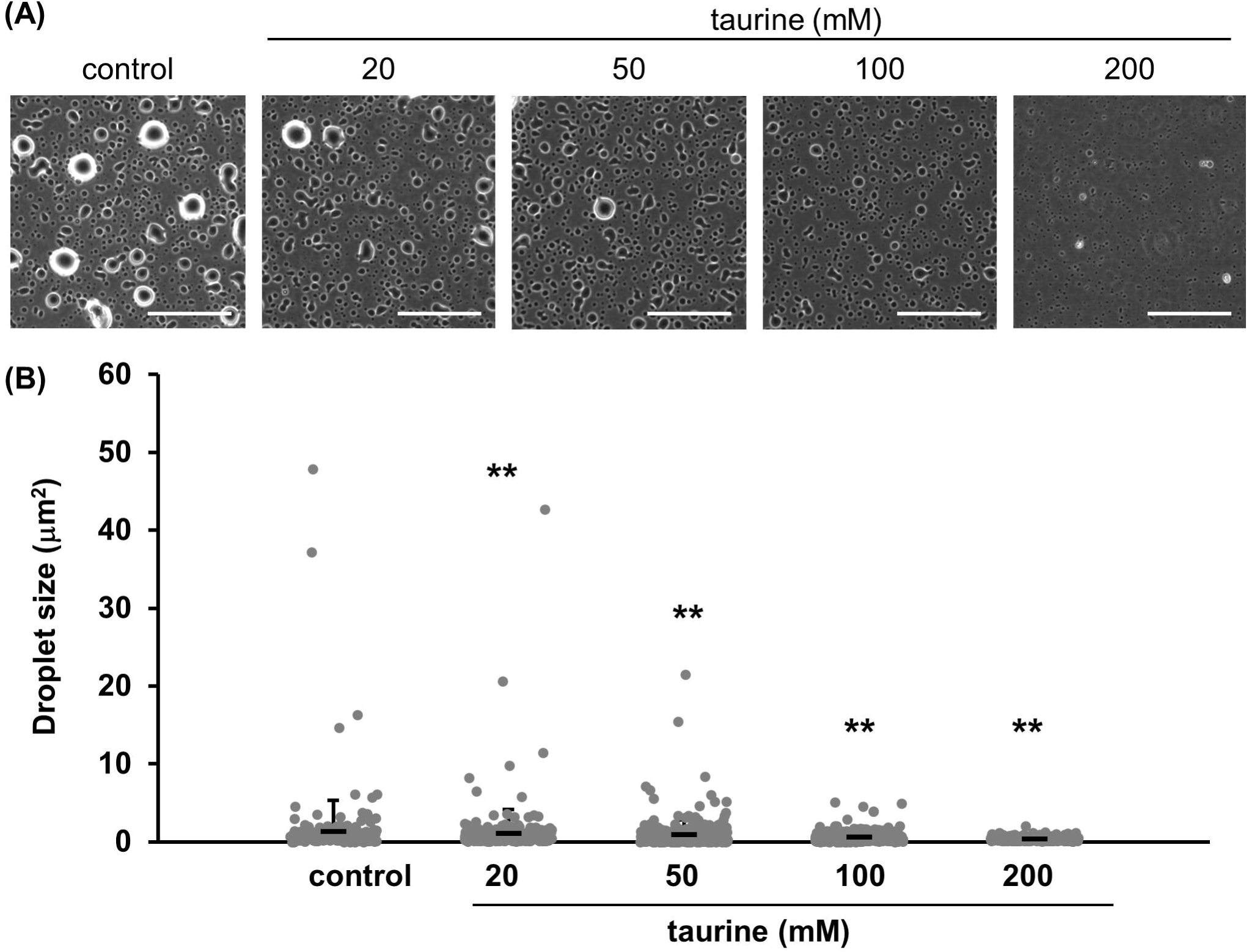
Effect of taurine on LLPS of lysozyme and ovalbumin mix solution (2.5 mg/mL each). (A) Optical microscopic images of heteroprotein mix solution with or without taurine (20∼200 mM). (B) Size of liquid droplets for heteroprotein solution with or without taurine. Experiments were performed in Tris buffer (5mM, pH 7.4). Gray circles indicate the size of each droplet. Mean droplet sizes with SD are indicated by black bars (n = 255-347). **; p<0.001 v.s. control. Scale bars = 100μm.

Furthermore, we confirmed LLPS formation of lysozyme and ovalbumin by using FITC-labelled lysozyme instead of non-labeled lysozyme to reveal whether lysozyme itself is contained in liquid droplets. As shown in Figure 4, liquid droplets were observed by fluorescent microscopy, indicating that FITC-labelled lysozyme forms LLPS. No fluorescent droplets were confirmed in heteroprotein mix solution without FITC-labeled lysozyme. Moreover, the addition of taurine decreased the number and size of liquid droplets.

**Figure 4.**
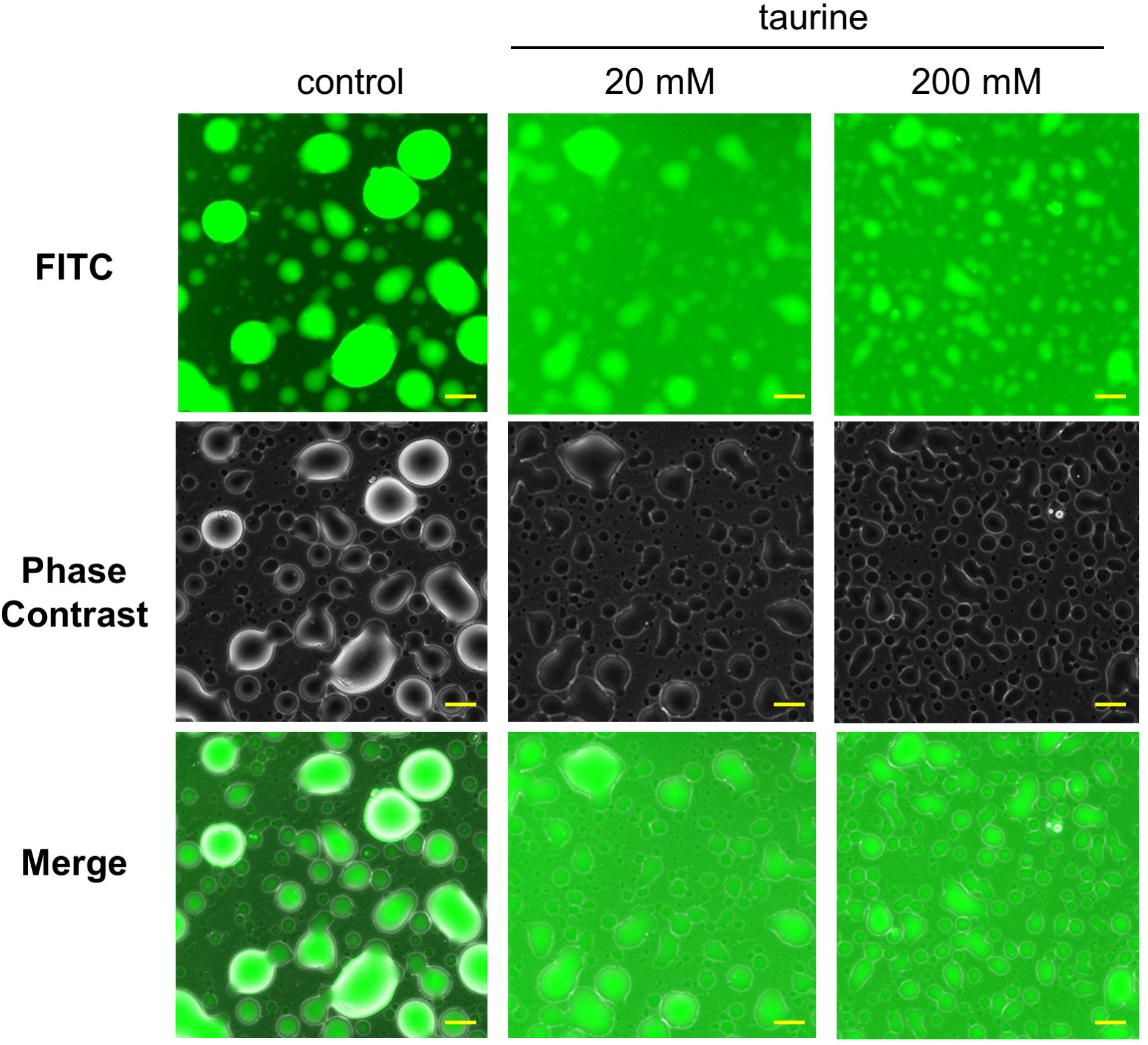
LLPS of FITC-labelled lysozyme in lysozyme and ovalbumin mix solution. Optical and fluorescent microscopic images of liquid droplet for FITC-lysozyme and ovalbumin mix solution in Tris buffer (5mM, pH 7.4) with or without taurine.

Since the previous reports used water as a reaction buffer instead of Tris buffer to form LLPS of lysozyme and ovalbumin, we also tested LLPS formation of two proteins in water (Figure S1). The addition of taurine did not largely change the pH of the heteroprotein solution. Increasing turbidity was observed in this condition, and the addition of taurine decreased turbidity. The droplet was confirmed by light microscopy. The addition of 200 mM taurine, but not 20 mM, inhibits LLPS formation.

## Discussion

In the present study, we demonstrated that taurine inhibits lysozyme-associated LLPS formation. Previous studies reported that taurine can weakly interact with lysozyme in solution. The surface of stable proteins contain positively and negatively charged amino acid side chains. Ionic interactions readily develop between these charged amino acid side chains and either salts or other charged compounds, including taurine. According to Abe et al taurine supports the folding of lysozyme under a crowding condition (Abe et al. 2015). Their NMR analysis revealed an interaction between taurine and hydrophilic residues. However, a cation-pi interaction of taurine with tryptophan residues has been reported. Meanwhile, beta-alanine, which is a similar chemical structure to taurine and has carboxylic acid instead of sulfonic acid, does not interact with lysozyme, indicating the specific action of taurine against lysozyme. Additionally, they also identified the interaction between taurine and the disordered regions of CedA protein and PriC N-terminal domain. Bruzdziak et al. demonstrated that the amino-group of taurine interacts with the side chain of lysozyme as assessed by thermodynamic analysis (Bruździak et al. 2018). Given the importance of the cation-#pai interaction and the disordered regions of proteins for LLPS formation, the weak interaction between taurine and proteins may contribute to the inhibition of LLPS formation.

There is also evidence that taurine alters water structure. This is not surprising because taurine is a zwitterion that has been found to be a water structure breaker. Both anions and cations that are water structure breakers have also been found to stabilize protein structure. Central to the action of the water structure breakers is the effect of nonpolar residues on water structuring. In the stable protein, nonpolar residues, such as isoleucine, are buried within the interior of the protein, which shields them from water. By contrast, in the denatured protein the nonpolar residues are exposed to aqueous medium. The exposed environment is thermodynamically unfavorable (entropy driven), as water forms large clusters around the nonpolar chains. Indeed, the most thermodynamically stable condition is one in which the nonpolar residues of proteins adhere to one another in the interior of the protein, so as to minimize water exposure, a condition called hydrophobic bonding. Some compounds compete with hydrophobic groups for available water molecules, thereby altering hydrophobic bonding. In the folded protein, the hydrophobic contribution is probably the most important stabilizing factor in proteins (Yancey 2005; Bruździak et al. 2018). The contribution of hydrophobic bonding has been demonstrated for the regulation of temperature-induced protein unfolding, urea denaturation and protein regeneration. The effect of taurine on water structure and hydrophobic interactions may also contribute to inhibition of LLPS formation, since protein-water interaction may be a driving force of LLPS formation (Ribeiro et al. 2019).

Meanwhile, the other osmolyte, trimethylamine oxide (TMAO), has the opposite effect of LLPS formers; the addition of TMAO to a solution containing #gamma-crystallin and TDP-43 protein enhances LLPS (Choi et al. 2018; Cinar et al. 2019). We also observed that TMAO increases LLPS formation of a heteroprotein mixture of lysozyme and ovalbumin (Figure S2). Therefore, the influence of osmolytes differ depending on each chemical.

Taurine functions as the cytoprotective metabolite through various mechanisms, such as antioxidation, Ca^2+^ handling control, and energy metabolism, etc. (Schaffer et al. 2010). Taurine also has diverse pharmacological properties in diseases, including heart failure, liver diseases, diabetes, and neurodegenerative disease (Ito et al. 2012, 2014; Menzie et al. 2014; Miyazaki and Matsuzaki 2014). The findings of the present study suggest that the action of taurine against LLPS formation may relate to its cytoprotective and pharmacological roles. For example, taurine has been reported to ameliorate neurodegenerative diseases in mice, such as Alzheimer’s disease model and Parkinson’s disease model (Kim et al. 2014; Che et al. 2018). Therefore, the inhibitory effect of taurine on fibril formation of amylogenic proteins, amyloid-#beta, and #alpha-synuclein may be associated with the prevention of LLPS formation. As another example, taurine prevents the hyperosmotic stress-induced formation of stress granules (Bounedjah et al. 2012). Taurine might directly interact with stress granule-related proteins to prevent assembly. Importantly, hyperosmotic stress induces the transcription of taurine transporter followed by an increase in cellular taurine concentration, which may lead to the dissolution of stress granules. Further studies are necessary to reveal the intracellular role of taurine against LLPS and taurine-related cellular functions.

## Acknowledgments

Hen egg lysozyme was kindly gifted from QP Corporation (Japan). This study was granted by Lotte Foundation (to Takashi Ito) and #a competitive grant from Fukui Prefecture University (to Takashi Ito).

